# Drosophila core circadian clock neurons peptidergically regulate activity of insulin-producing cells

**DOI:** 10.64898/2026.03.23.713581

**Authors:** Naureen A. Hameed, Sergio L. Crespo Flores, Evan Cirone, Chenyue Zhao, Annika F. Barber

## Abstract

Central pacemaker neurons use a combination of external stimuli and neuropeptide signaling to synchronize molecular oscillations leading to circadian behaviors. The clock network structure and signaling between these pacemaker neuron groups have been well described, but how these pacemakers communicate with specific brain output regions remains poorly understood. Here, we identified how “core” clock neurons in *Drosophila*, the ventrolateral neurons (LNvs), signal to the proto-hypothalamic region, the *pars intercerebralis* (PI). We provide evidence to show that LNvs functionally modulate insulin-producing cells (IPCs) of the PI in a time-of-day-dependent manner. This functional connectivity relies on neuropeptidergic signaling of two classical clock neuropeptides: pigment dispersing factor (PDF) and short Neuropeptide F (sNPF). Connectomic analysis does not identify any direct synaptic inputs from clock neurons to IPCs. Small ventrolateral clock neurons, which secrete both PDF and sNPF are 15-20 mm away from IPCs, suggesting that volume transmission across these distances may be possible in the fly dorsal protocerebrum. Peptide application with functional imaging of IPCs provides insight into how these two neuropeptides may act synergistically via their receptors to signal to IPCs. Our findings indicate that LNvs can signal to IPCs by volume transmission and also form indirect multisynaptic circuits with IPCs. Thus, neuropeptides associated with intra-clock signaling also serve as circadian cues to non-clock neurons, which may model more broadly how clock peptides communicate to clock output regions.

**Author Summary:** Circadian clocks in the brains of animals from flies to humans allow animals to anticipate daily environmental changes and temporally coordinate internal processes. The central clock in the brain serves as a master pacemaker that sets the pace of circadian clocks in all other tissues. Fruit flies have been an essential model system for discovering the genetic and cellular underpinnings of circadian rhythms, with work in flies being awarded the 2017 Nobel Prize in Physiology or Medicine. Brain clocks in flies and mammals use classical synaptic communication and neuromodulatory peptides to signal between the neurons that make up the brain clock. Our study asks whether the neuropeptides that signal between clock neurons also signal from clock to non-clock neurons. We found that the signature fly clock neuropeptide pigment dispersing factor signals outside the brain clock to insulin producing cells of the brain. There are no synaptic connections from clock neurons to insulin producing cells, instead signaling occurs by diffusion of peptide signals across tens of microns, termed volume transmission. Our findings point to the possibility that neuropeptide volume transmission may be a general feature not only of intra-clock signaling but also of brain clock output signaling to non-clock neurons.

## Introduction

Circadian behavior and physiology are regulated by a central brain pacemaker that provides entrainment signals to the molecular clocks of downstream cells and tissues. Across species, the brain clock is structured as a network of molecular oscillators that together serve to synchronize circadian oscillation within the brain clock and provide robust outputs [1,2]. Work in model organisms has served to elucidate the brain clock network structure, while less is known about how these multiple oscillators coordinate signals to output regions.

The *Drosophila* brain clock network consists of ∼240 neurons that express the cell-autonomous clock transcription-translation feedback loop [3–5]. Although fly clock neurons are distributed throughout the central brain, rather than in a single nucleus as in mammals, the essential clock circuit structure is conserved from flies to humans [5–7]. Neuropeptides are crucial signaling molecules within brain clocks. In flies, the signature clock neuropeptide is pigment dispersing factor (PDF) which is produced by the small and large ventrolateral neurons (s- and l-LNvs), except for the 5^th^ s-LNv which is PDF negative [8]. PDF serves to coordinate clock neuron groups by adjusting the period and phase of the molecular clock and regulating cellular excitability [9–13]. s-LNvs co-express short neuropeptide F (sNPF) and glycine, while a subset of l-LNvs co-express neuropeptide F (NPF) [14–16]. s-LNvs are morning active, and release neuropeptides into the dorsal protocerebrum, signaling to dorsolateral (LNd) and dorsal clock neurons (DN) [11,17–19]. LNd neurons are evening active and consist of four distinct subgroups identifiable by transcriptomic and connectomic differences [16,19,20].

Notably, only the six CRYPTOCHROME-expressing (CRY^+^) LNds express PDF receptor (PDFR) [21]. Four of the CRY^+^ LNds signal via sNPF and acetylcholine, while two signal via NPF and ion transport peptide [8]. Dorsal neurons are divided into three major groups (DN1-3) and neurons within all groups are PDF-responsive [22–24]. The dorsal neurons express diverse clock neuropeptides, including sNPF expression in the four DN2s and some of the 28 night-active posterior DN1s (DN1p) [5,15,16,19].

Clock neurons signal not only within but also outside of the clock network, including to neurosecretory hubs in the lateral horn and the *pars intercerebralis* (PI) [25]. The PI is a major clock output region implicated in regulating circadian locomotor and feeding rhythms and peripheral gene expression rhythms [26–32]. Identified PI neuron populations express multiple neuropeptides including the *Drosophila* insulin-like peptides (DILPs) 2, 3, and 5, and diuretic hormone 44 (DH44) [27,31,33–36]. Although PI neurons lack molecular clock gene expression, they display rhythmic firing patterns and intracellular calcium levels entrained by brain clock neurons [28,37]. Circuit tracing methods have identified signaling from LNd and DN1 clock neurons to insulin-producing cells (IPCs) and *DH44*^+^ PI neurons of the PI [27–30]. Single-cell RNAseq datasets, antibody labeling, and *in situ* hybridization experiments suggest that some IPCs and *SIFa*+ neurons co-express PDFR and sNPFR, while *DH44*^+^ neurons express only sNPFR [38–41]. IPCs respond to application of synthetic PDF and sNPF peptides with increased cyclic AMP levels, confirming functional roles for receptor expression [42]. Although the fly brain connectome does not identify direct synaptic connections from any clock neuron to the PI, proximity analysis demonstrates that s-LNv neurons come within 15 microns of IPCs, allowing for the possibility of peptidergic volume transmission [5,43,44]. We hypothesized that LNv clock neurons provide input to PI neurons via neuropeptide signals. We provide evidence that PDF^+^ LNvs provide morning-specific input to IPCs, but not *DH44*^+^ neurons. CRISPR knockdown of PDFR and sNPFR confirmed prior findings that loss of PDFR blunts the IPC response to sNPF, and further showed that loss of sNPFR completely abrogates the IPC response to PDF. Our data support a model in which LNvs activate some IPCs both directly via bulk neurotransmission and others indirectly via sNPF from other PDF-responsive clock neurons.

## Results

### Functional imaging identifies time-of-day dependent modulation of IPCs, but not *DH44*^+^ neurons, by LNv stimulation

Based on prior findings suggesting peptidergic modulation of IPCs by LNv neuropeptides and connectomics suggesting proximity between s-LNvs and IPCs, we sought to confirm functional connectivity using a GCaMP-based stimulus-response assay in acutely dissected brains (Fig. S1) [5,28,29,42,45]. In this assay, the ATP-gated cation channel P2X2 is expressed in the putative upstream LNvs and the genetically encoded calcium indicator GCaMP6m was expressed in PI neurons to allow monitoring of intracellular calcium levels before and after clock neuron stimulation by application of 2.5 mM ATP (Fig 1, green symbols) [28,45,46]. As a genetic control for leaky expression of the P2X2 channel, we expressed GCaMP6m in PI neurons of flies expressing UAS-P2X2 without *PDF-Gal4*-driven expression and quantified the change in GCaMP fluorescence relative to average pre-ATP application baseline signal (ΔF/F). We did not observe a significant change in IPC GCaMP signal upon ATP application in genetic controls (Fig 1, grey symbols). In the morning (ZT 0-4), we observed an increase in GCaMP6m fluorescence intensity in male IPCs when P2X2-expressing LNv clock neurons were stimulated by ATP application (ΔF/F = 1.54 ± 0.16), a significant increase compared to genetic negative control brains (ΔF/F = 1.04 ± 0.16) (Fig 1A, 1A’). IPC activation by LNv stimulation is heterogeneous, with some cells showing large increases in GCaMP6m signal while other cells in the same brain exhibited little to no response. This activation is diurnal, as stimulating LNv neurons in the evening (ZT 11-15) did not result in significant GCaMP6m activation in IPCs in male brains (Fig 1B, 1B’). Based on recent findings of sexually dimorphic roles for PDF in circadian timekeeping, we also tested LNv to IPC functional connectivity in female brains [47]. Female brains showed a similar morning-specific mean increase in IPC GCaMP6m fluorescence in response to LNv neuron stimulation (ΔF/F = 1.34 ± 0.09). Female flies had similar within-brain heterogeneity of response as males, though both the average and maximal GCaMP6m responses to LNv stimulation were smaller than in male flies (Fig 1C, 1C’). As in males, in the evening (ZT 11-15), female brains show no IPC GCaMP response to LNv neuron stimulation (ΔF/F = 1.14 ± 0.06) (Fig 1D-D’).

**Fig 1.**
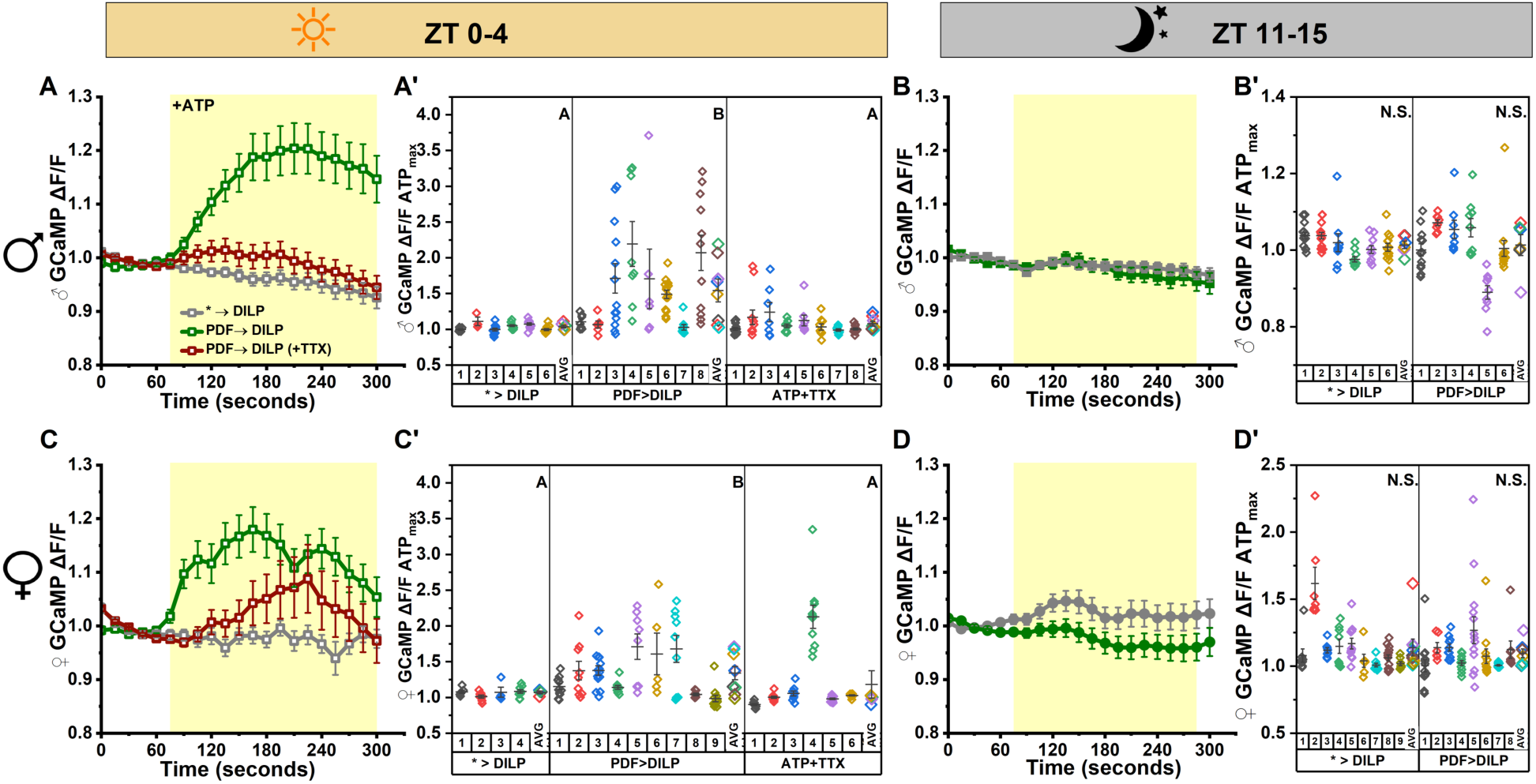
LNv neurons functionally modulate IPCs neurons in a time-of-day dependent manner. (A) GCaMP6m signal over time in *DILP2*^+^ neurons during PDF neuron activation (green, n= 8 brains, 73 cells) imaged at ZT 0-4 in male brains. Genetic controls lacked LexAop-P2X2 (gray, n=6 brains, 34 cells). Sodium channel blocker TTX was applied together with ATP in brains indicated (red, n=8 brains, 78 cells) ATP or ATP+TTX application is indicated by yellow shading. (A’) Maximum change in GCaMP intensity during activation for each cell by brain and brain averages by genetic line/treatment from panel A. Each small point represents an individual cell for each brain, while larger symbols in the right column of each line/treatment panel indicate the mean response per brain matching individual symbol colors by brain. (B) Plotted as in panel A for ZT 11-15 (experimental: n=6 brains, 60 cells, control: n=6 brains, 70 cells). (B’) Plotted as in panel A’ for panel B. (C) Plotted as in panel A in female brains (experimental: n=9 brains, 83 cells, control: n=4 brains, 28 cells, TTX+ATP: n=6 brains, 63 cells). (C’) Plotted as in panel A’ for panel C. (D) Plotted as in panel A imaged at ZT 11-15 for female brains (experimental: n= 8 brains, 84 cells, control: n=9 brains, 79 cells) (D’) Plotted as in panel A’ for panel D. Data are presented as mean ± SEM for cells. Brain means that do not share a letter are significantly different; N.S. indicates no significant difference. One-way ANOVA analysis with Tukey’s multiple comparisons done across genetic lines/ treatment groups using brain-averaged responses for all panels except for non-normal ATP-TTX data in panel C’ (Kruskal-Wallis ANOVA with Dunn’s test). *P ≤* 0.05

To test whether the morning LNv to IPC connection is monosynaptic, we repeated the stimulus-response experiment in the presence of the voltage gated sodium channel blocker tetrodotoxin (TTX). TTX application prevents indirect activation of IPCs by other LNv targets, while ATP stimulation of P2X2 in LNvs should provide the depolarizing stimulus to permit neurotransmitter and neuropeptide vesicle release, allowing us to test the direct connection. In males, co-application of ATP and 2 μM TTX reduced the GCaMP IPC response to LNv stimulation to ΔF/F = 1.07 ± 0.03, which was not significantly different from genetic controls (Fig 1A, maroon). These data suggest that most morning excitatory LNv input to IPCs occurs via indirect, action potential-dependent signaling through LNv targets that serve as intermediaries in the LNv to IPC circuit. In females, co-application of ATP and 2 μM TTX also reduced the mean IPC response to LNv stimulation to ΔF/F = 1.18 ± 1.15, which was not significantly different from genetic controls (Fig 1C, maroon). While the mean IPC response to LNv stimulation in the presence of TTX is not significantly different from negative controls in either sex, it should be noted that one or more IPCs respond in approximately half of sample brains, suggesting the possibility that a few IPCs may be sensitive to direct LNv signaling.

The LNd and DN1 clock neurons provide functional input to *DH44+* in addition to IPCs. To test whether LNvs similarly provide input to diverse PI cell groups, we also tested the functional connectivity between LNv clock neurons and *DH44*+ PI neurons using the same stimulus-response assay. However, stimulation of P2X2 expressing LNv neurons by ATP application did not elicit a significant change in mean GCaMP response in *DH44*+ neurons compared to genetic negative controls at ZT 0-4 or ZT 11-15 in male (Fig 2A, 2B) or female (Fig 2C, 2D) brains. Females did, however, show a slow increase in GCaMP6m signal that persisted beyond the ATP application window in the morning (Fig 2C, 2C’).

**Fig 2.**
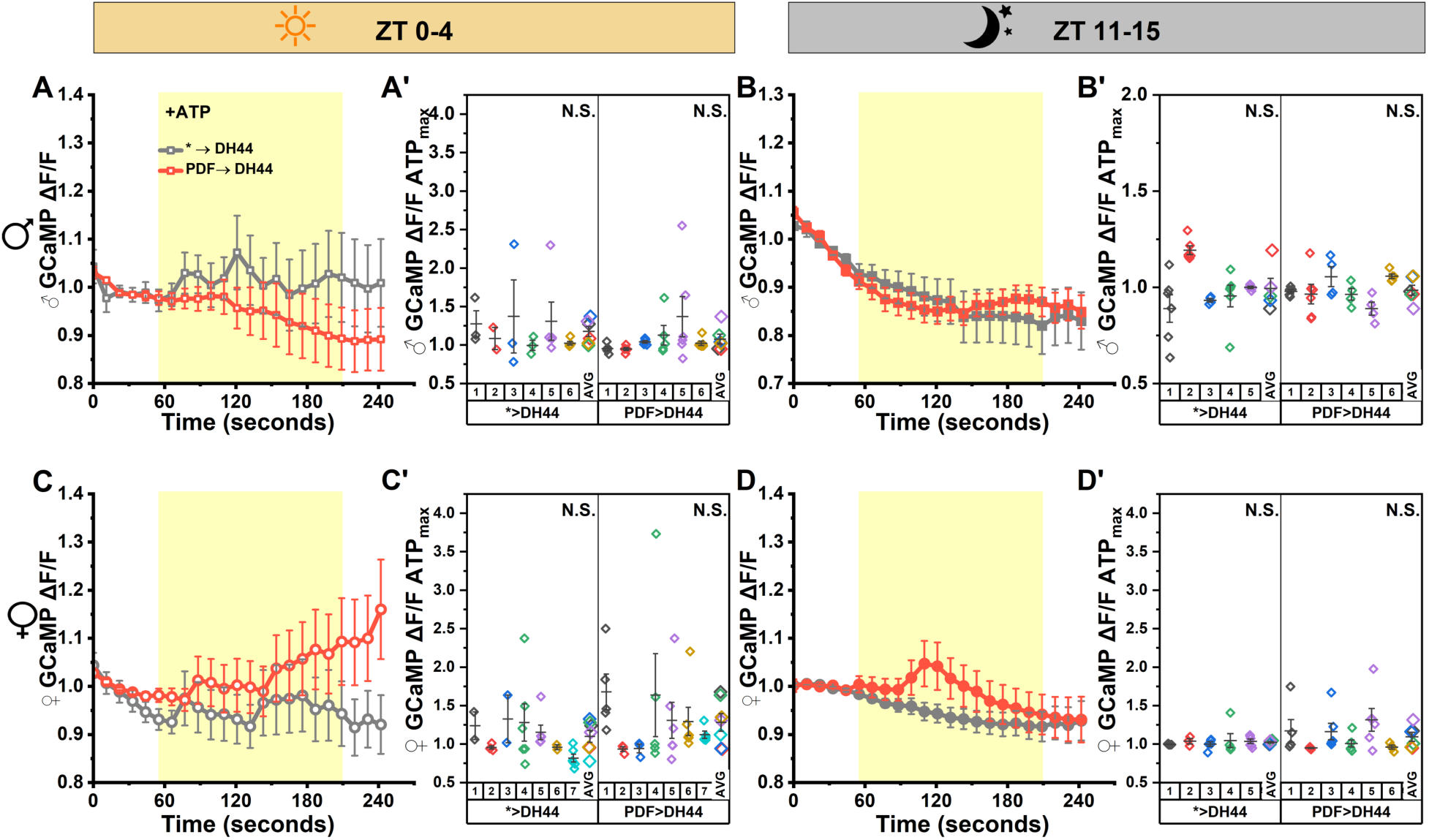
LNv neurons do not functionally modulate DH44+ neurons. (A) GCaMP6m signal over time in *DH44^+^* neurons during PDF neuron activation (orange, n= 6 brains, 21 cells) imaged at ZT 0-4 in male brains. Controls (gray, n=6 brains, 33 cells) lacked LexAop-P2X2. ATP application is indicated by yellow area. (A’) Maximum change in GCaMP intensity during activation for each cell by brain and brain averages by genetic line from panel A. Each small point represents an individual cell for each brain, while larger symbols at the right of the panel for each genetic line indicate the mean response per brain matching individual symbol colors by brain (B) Plotted as in panel A imaged at ZT 11-15 (experimental: n=5 brains, 27 cells, control: n=6 brains, 27 cells) (B’) Plotted as in panel A’ for panel B. (C) Plotted as in panel A in female brains (experimental: n=7 brains, 27 cells, control: n=7 brains, 33 cells) (C’) Plotted as in panel A’ for panel C. (D) Plotted as in panel A imaged at ZT 11-15 for female brains (experimental: n= 5 brains, 25 cells, control: n=6 brains, 31 cells). (D’) Plotted as in panel A’ for panel D. Data are presented as mean ± SEM for cells. Brain means that do not share a letter are significantly different; N.S. indicates no significant difference. One-way ANOVA analysis with Tukey’s multiple comparisons done across genetic lines. *P ≤* 0.05

### PDF neuropeptide activation of IPCs is time-of-day dependent and requires both PDFR and sNPFR

Large and small LNv clock neurons signal via the neuropeptide PDF, while s-LNvs also express sNPF and glycine [8,14,15,48]. PDF is often seen as the signature circadian cue in flies, while sNPF is more broadly expressed. These neuronal signals have well-defined roles in LNv neuron communication to downstream neurons within the clock network, but our functional imaging data suggested that LNv neurons may also signal directly to non-clock IPCs via their classical neuropeptides. *Drosophila* IPCs express sNPFR in some IPCs of the adult brain and respond to application of both PDF and sNPF exogenous peptides by increasing cAMP levels, suggesting PDFR may also be present [40–42,49]. Imaging studies show that s-LNv clock neuron dendrites are in close proximity to fine IPC processes when imaged at ZT 1, a time that s-LNv dorsal projections are maximally extended [5,50,51]. We confirmed that morning (ZT 0-4) bath application of 10 μM PDF peptide results in increased IPC GCaMP signal in male brains (ΔF/F = 1.29 ± 0.06) and female brains (ΔF/F = 1.18 ± 0.04) at ZT 0-4 compared to vehicle control (Fig 3A, 3C, grey symbols). The IPC response to PDF peptide application was morning specific. In both sexes PDF peptide application at ZT 11-15 did not increase GCaMP ΔF/F (Fig 3B, B’, D, D’).

**Fig 3.**
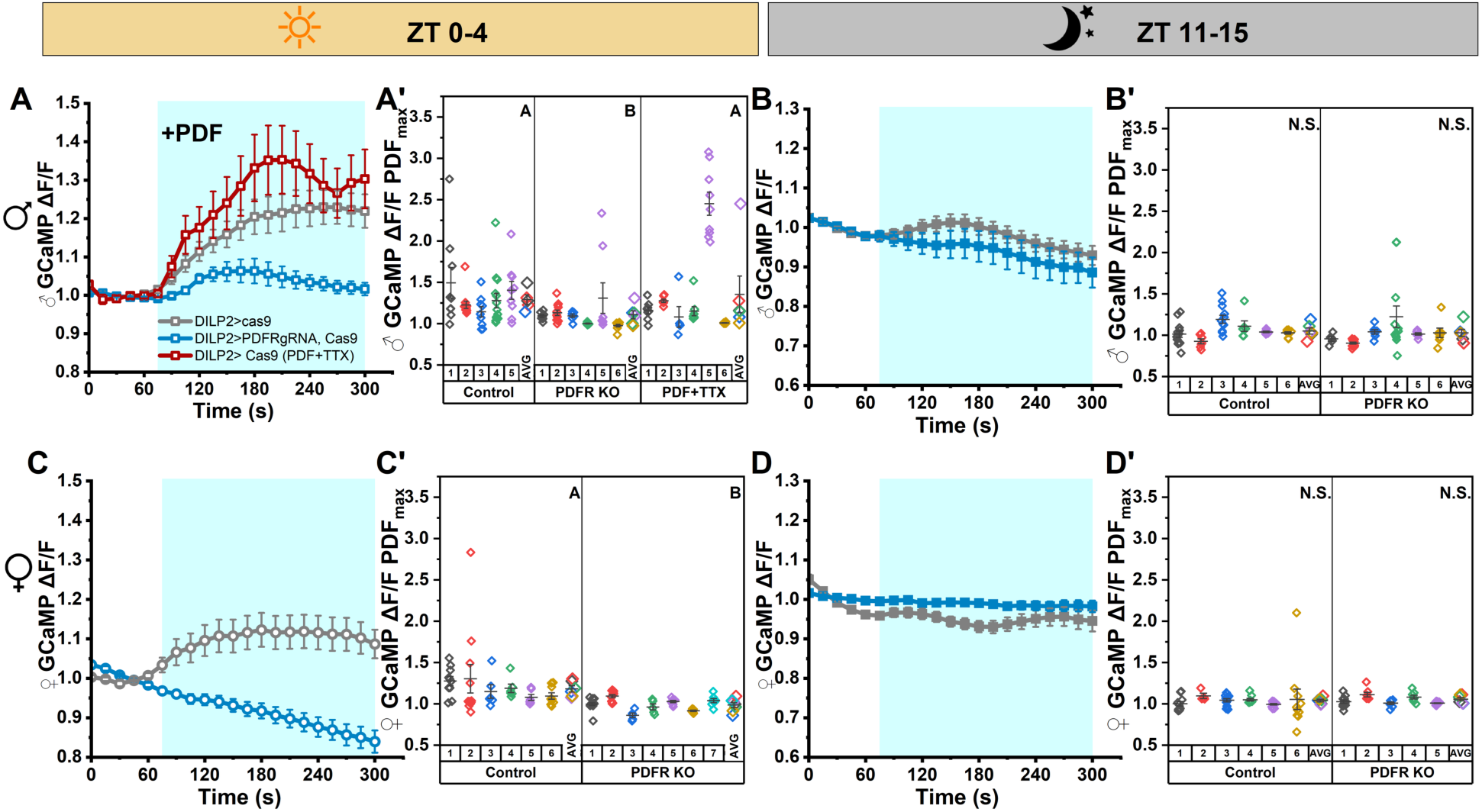
PDF neuropeptide functionally modulates IPCs in a time-of-day dependent manner directly via PDF receptor. (A) GCaMP6m signal over time in IPCs with (blue: n=6 brains, 52 cells) and without (gray: n=5 brains, 53 cells) PDFR CRISPR knockout from male brains imaged at ZT 0-4 during perfusion of artificial hemolymph, followed by perfusion of 10 μM PDF. Sodium channel blocker TTX was applied along with PDF in brains indicated (red, n=6 brains, 40 cells). Blue area indicates timing of PDF application on brains. Data are represented as mean ± SEM. (A’) Maximum change in GCaMP intensity during activation for each cell by brain and brain averages by genetic line from panel A. Each small point represents an individual cell for each brain, while larger symbols at the right of the panel for each genetic line indicate the mean response per brain matching individual symbol colors by brain. (B) Plotted as in panel A imaged at ZT 11-15 (blue: n=6 brains, 51 cells, gray: n= 6 brains, 54 cells). (B’) Plotted as in panel A’ for panel B. (C) Plotted as in panel A from female brains (blue: n=7 brains, 57 cells, gray: n=6 brains, 48 cells). (C’) Plotted as in panel A’ for panel C. (D) Plotted as in panel A from female brains imaged at ZT 11-15 (blue: n=5 brains, 46 cells, gray: n= 6 brains, 58 cells). (D’) Plotted as in panel A’ for panel D. Data are presented as mean ± SEM for cells. Brain means that do not share a letter are significantly different; N.S. indicates no significant difference. One-way ANOVA analysis with Tukey’s multiple comparisons done across genetic lines except for non-normal ATP-TTX data in panel A’ (Kruskal-Wallis ANOVA analysis with Dunn’s test). *P ≤* 0.05.

To test the role of PDFR in the IPC response to LNv clock neuron signaling, we created CRISPR tools for cell-specific disruption of the *pdfr* gene, based on previous work [52–55]. We then used peptide application to confirm the efficacy of our IPC-specific PDFR CRISPR knockout, which showed significantly blunted IPC response to PDF application in the morning in both male and female flies (Fig 3A, 3C, blue symbols). In males, the morning response was still present, but with a significantly reduced amplitude compared to genetic parental controls (ΔF/F = 1.11 ± 0.05) (Fig 3A, 3A’). To validate that PDF application regulates IPC GCaMP6m activation directly, we co-applied PDF with TTX and observed an increase in GCaMP response (ΔF/F = 1.30 ± 0.19). There was no significant difference in GCaMP ΔF/F between PDF+TTX and PDF application alone, aligned with prior findings from Nagy *et al*. using a cyclic AMP sensor and demonstrating that PDF can activate IPCs in the absence of other PDF-active synaptic inputs. Female brains also showed no IPC GCaMP response to PDF application in the absence of PDFR (ΔF/F = 1.00 ± 0.03, Fig 3C, 3C’). There is no response to PDF application in the evening in male or female flies after PDFR CRISPR (Fig 3B, 3D). These results provide further support for the capacity for IPCs to respond to PDF and show a role for PDFR in regulating the magnitude and timing of the calcium response to PDF.

sNPF is broadly expressed in the fly brain, and in the clock circuit it is expressed by the s-LNvs and two CRY^+^ LNd clock neurons. sNPF regulation of IPCs in larval and adult stages affects growth and metabolism, and IPCs respond to sNPF application with increases in cAMP [40,42,49]. The effect of sNPFR activation is cell type-specific, with examples of both inhibition and activation [56,57]. We found that IPCs also respond to sNPF application with increases in intracellular calcium, as measured by GCaMP6M intensity, and this response is morning specific. Morning (ZT 0-4) bath application of 10 μM sNPF peptide results in increased IPC GCaMP6m signal compared to baseline perfusion, with an increase in male (ΔF/F = 1.62 ± 0.21) and female brains (ΔF/F = 1.18 ± 0.04) at ZT 0-4 (Fig 4A, A’, C, C’, grey symbols). The IPC response to sNPF peptide application was morning specific, as we found no change in GCaMP6m signal in response to sNPF peptide application at ZT 11-15 compared to the pre-sNPF vehicle application baseline in male or female flies (Fig 4B, B’, D, D’). CRISPR knockdown of sNPF receptor resulted in a total loss of GCaMP6m response to sNPF application (ΔF/F = 1.06 ± 0.01) in the morning (ZT 0-4), which was significantly lower than the response to sNPF in wild type IPCs (Fig 4A, A’, C, C, purple symbols) In the evening, when IPCs do not respond to sNPF application, there was no significant difference in GCaMP6m response from vehicle control baseline or from positive control flies with sNPFR CRISPR (Fig 4B, B’, D, D’).

**Fig 4.**
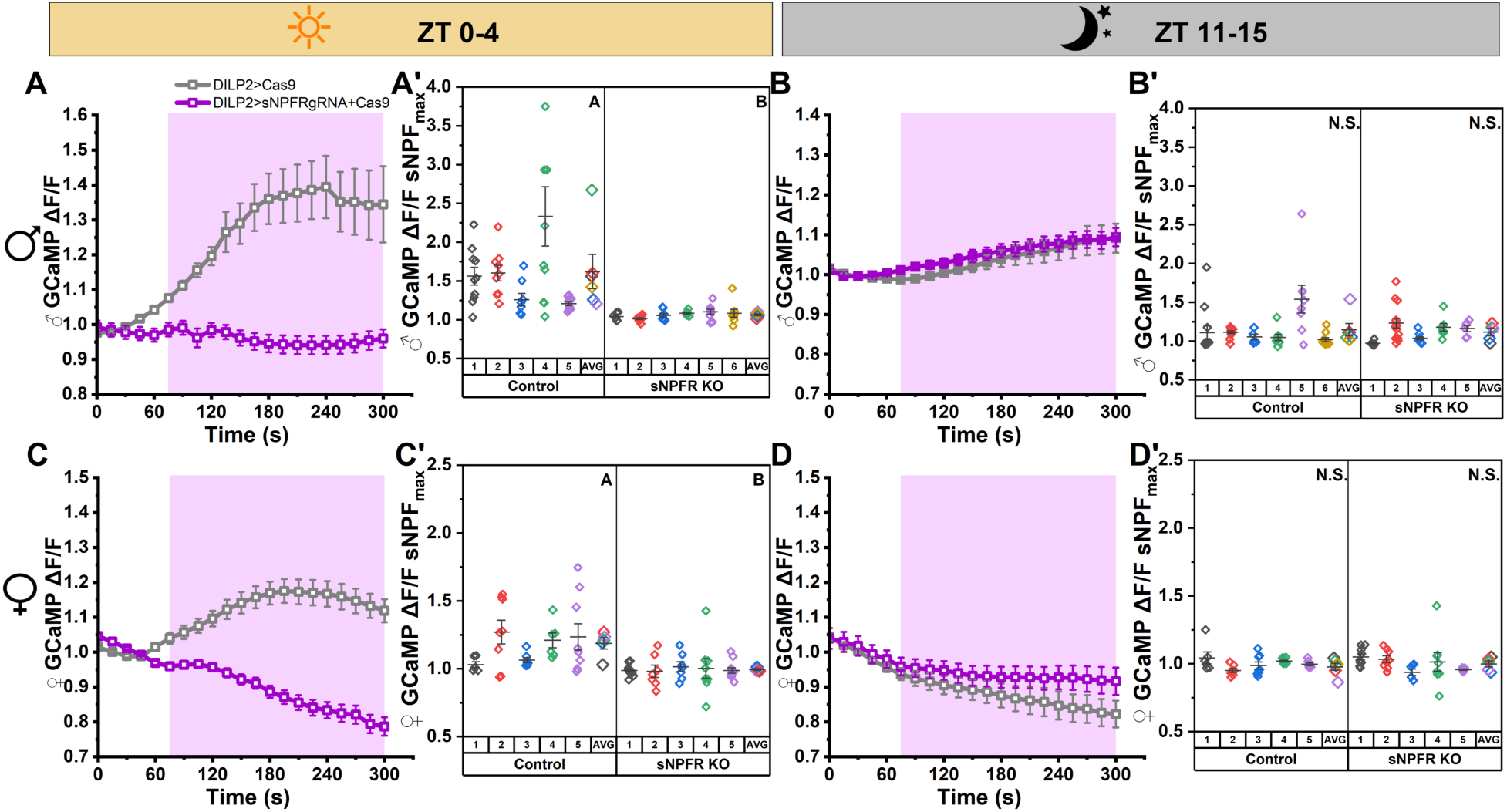
sNPF neuropeptide functionally modulates IPCs in a time-of-day dependent manner directly via sNPF Receptor. (A) GCaMP6m signal over time in IPCs with (magenta: n=6 brains, 47 cells) and without (gray: n=5 brains, 47 cells) sNPFR CRISPR knockout from male brains imaged at ZT 0-4 during perfusion of artificial hemolymph, followed by perfusion of 10 μM sNPF. Magenta area indicates timing of sNPF application on brains. Data are represented as mean ± SEM. (A’) Maximum change in GCaMP intensity during activation for each cell by brain and brain averages by genetic line from panel A. Each small point represents an individual cell for each brain, while larger symbols at the right of the panel for each genetic line indicate the mean response per brain matching individual symbol colors by brain (B) Plotted as in panel A imaged at ZT 11-15 (magenta: n=5 brains, 45 cells, gray: n= 6 brains, 57 cells). (B’) Plotted as in panel A’ for panel B. (C) Plotted as in panel A from female brains (magenta: n=5 brains, 46 cells, gray: n=5 brains, 44 cells) (C’) Plotted as in panel A’ for panel C. (D) Plotted as in panel A from female brains imaged at ZT 11-15 (magenta: n=5 brains, 38 cells, gray: n=5 brains, 38 cells). (D’) Plotted as in panel B for panel G. Data are presented as mean ± SEM for cells. Brain means that do not share a letter are significantly different; N.S. indicates no significant difference. Kruskal-Wallis analysis with Dunn’s multiple comparisons done across genetic lines for all panels. *P ≤* 0.05

In agreement with prior findings [42], we found that when PDFR was knocked out in IPCs, the morning-specific response to sNPF application was also blunted (Fig. 5A, 5B, cyan symbols). In the morning, male and female flies exhibited a small GCaMP6m response to sNPF application (ΔF/F_male_ = 1.38 ± 0.07, ΔF/F_female_ 1.30 ± 0.08,, Fig 5A, 5A’, 5B, 5B’). Expanding on prior findings, we were also able to test the response of IPC-specific sNPFR knockout to PDF application. In the morning, a time when IPCs are responsive to sNPF application (grey symbols), neither male nor female brains showed a GCaMP6m response to PDF application in the absence of sNPFR (ΔF/F_male_ = 1.23 ± 0.12, ΔF/F_female_ = 1.00 ± 0.01, Fig. 5C, C’, D, D’, purple symbols). At night, when IPCs normally do not respond to sNPF or PDF application, IPC-specific PDFR CRISPR did not have a significant GCaMP6m response to sNPF application compared to vehicle control baseline in either sex (Fig S2A, A’, B, B’). Similarly, at night, IPC-specific sNPFR CRISPR did not have a significant GCaMP6m response to PDF application compared to vehicle control baseline in either sex (Fig S2C, C’, D, D’). Although these experiments were performed with only a small number of animals, we did not further follow up on at this non-responsive time point.

**Figure 5.**
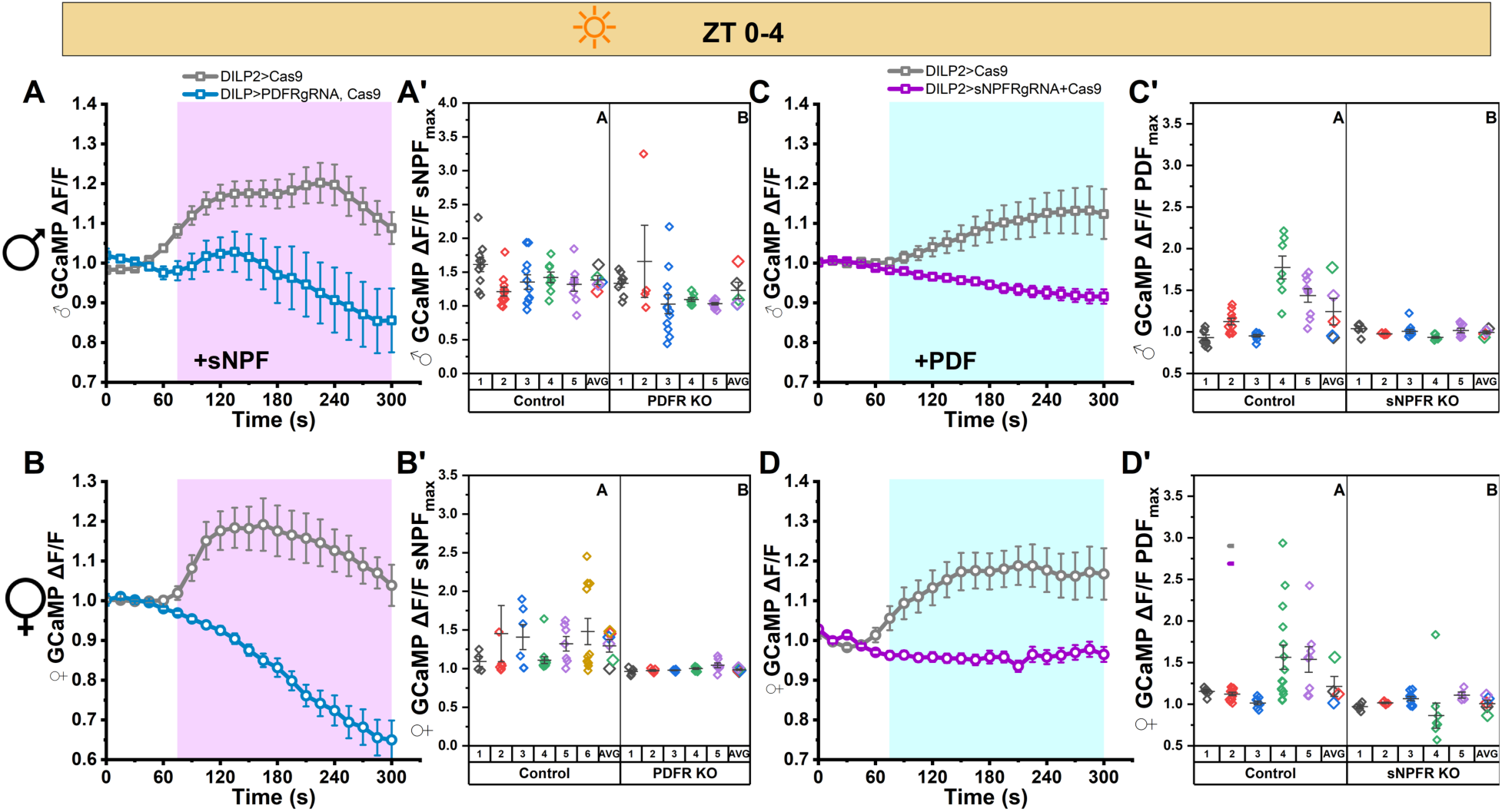
PDF and sNPF require reciprocal receptor function to activate IPCs. (A) GCaMP6m signal over time in IPCs with (blue: n=5 brains, 43 cells) and without (gray: n=5 brains, 49 cells) PDFR knockout from male brains imaged at ZT 0-4 during perfusion of artificial hemolymph, followed by perfusion of 10 μM sNPF. Magenta area indicates timing of sNPF application on brains. (A’) Maximum change in GCaMP intensity during activation for each cell by brain and brain averages by genetic line from panel A. Each small point represents an individual cell for each brain, while larger symbols at the right of each line panel indicate the mean response per brain matching individual symbol colors by brain. (B) Plotted as in panel A from female brains (blue: n=5 brains, 31 cells, gray: n=6 brains, 48 cells) (B’) Plotted as in panel A’ for panel B. (C) GCaMP6m signal over time in IPCs with (magenta: n=5 brains, 42 cells) and without (gray: n=5 brains, 44 cells) sNPFR knockout from male brains imaged at ZT 0-4 during perfusion of artificial hemolymph, followed by perfusion of 10 μM PDF. Blue area indicates timing of PDF application on brains. Data are represented as mean ± SEM. (C’) Maximum change in GCaMP intensity during PDF application plotted as in panel A’ for C. (D) Plotted as in panel C from female brains (blue: n=5 brains, 31 cells, gray: n=5 brains, 50 cells) (D’) Plotted as in panel C’ for panel C. Data are represented as mean ± SEM. *P<0.05, One-way ANOVA analysis done across genetic lines using brain-averaged responses. Data are presented as mean ± SEM for cells. Brain means that do not share a letter are significantly different; N.S. indicates no significant difference. One-way ANOVA analysis with Tukey’s multiple comparisons done across genetic lines. *P ≤* 0.05

### Connectomic analysis identifies proximity but no synaptic connections from clock neurons to IPCs

While our functional imaging data indicate the possibility of a weak synaptic connection from LNvs to IPCs, neuropeptide signaling by bulk neurotransmission can occur with close proximity in the absence of a synapse [58,59]. We used the female adult fly brain (FAFB) and male CNS (MCNS) connectomes to examine the proximity of PDF^+^ and sNPF^+^ clock neurons to IPCs [43,44]. We first screened for PDF^+^ or sNPF^+^ clock neurons within 15 μm of any IPC based on a previously-estimated distance for PDF signaling of 15 μm [5]. In the female connectome, this analysis identified a single left hemisphere s-LNv, as previously described, as well as the CRY^+^ LNds and the 5^th^ s-LNv clock neuron in both hemispheres. In the male connectome, proximity analysis again identified the CRY^+^ LNds and the 5^th^ s-LNv clock neuron in both hemispheres within 15 μm of IPCs. In the male connectome, there are no s-LNvs in either hemisphere within 15 μm of any IPC, expanding the proximity threshold to 20 μm identified a single right hemisphere s-LNv (Fig S3). In both male and female connectomes, l-LNvs do not come within 15 μm of IPCs.

We then re-screened the CRY^+^ LNds and the 5^th^ s-LNv clock neuron for contacts within 1 μm of any IPC. For the 5^th^ s-LNv, in the male connectome we found 14 contacts within 1 μm of IPCs (Fig 6A, A’), while in the female connectome we found 4 contacts (Fig 6B, B’). For the CRY^+^ LNds, in the male connectome we found 27 contacts within 1 μm of IPCs (Fig 6C, C’), while in the female connectome we found only 5 contacts (Fig 6D, D’). While each connectome represents a single brain at a single timepoint and many factors affect the likelihood of volume transmission, these findings suggest the possibility of sexually dimorphic sNPF volume transmission from LNds to IPCs.

**Fig 6.**
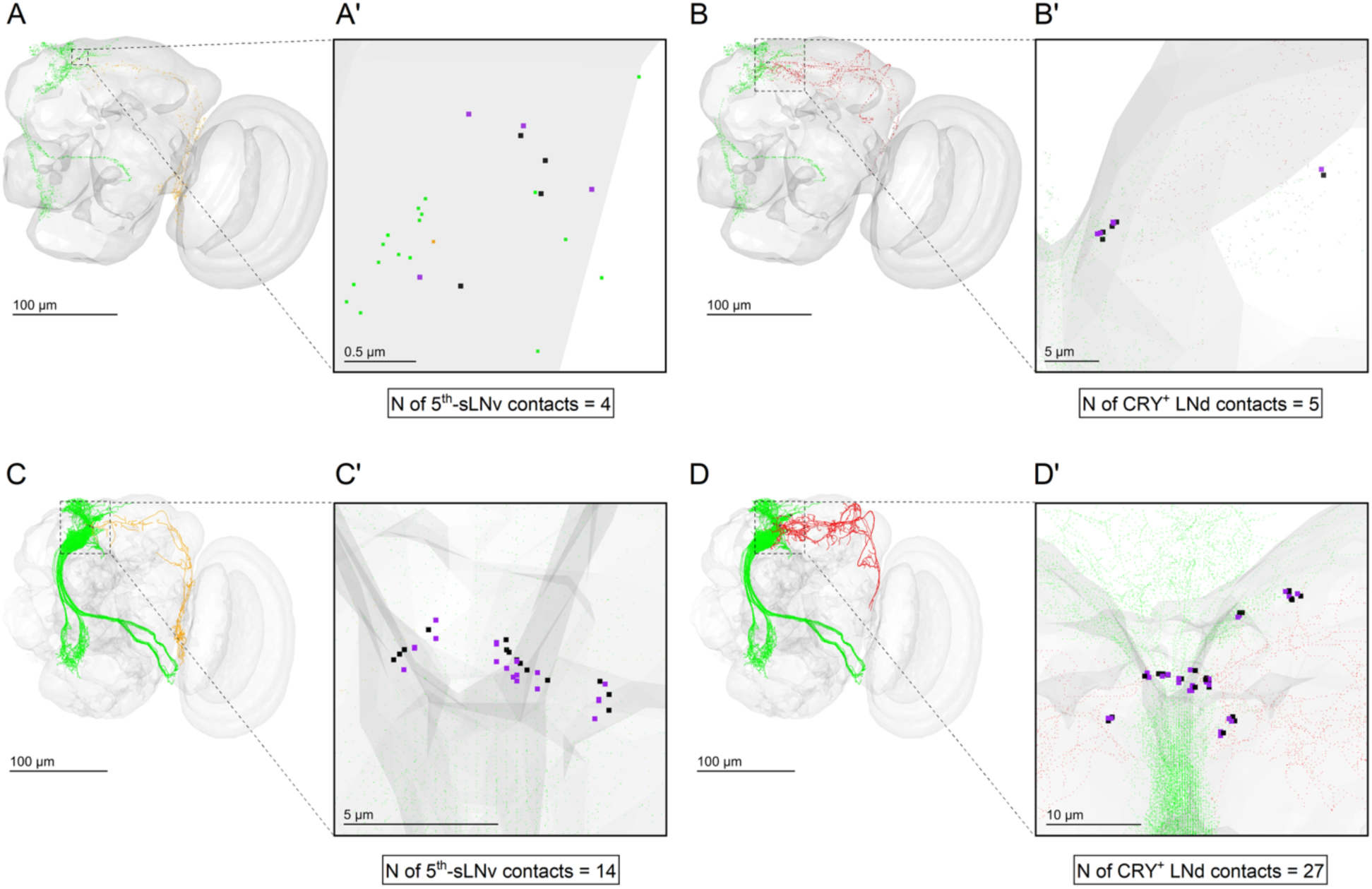
IPCs come within 1 µm proximity with the 5^th^ s-LNv and CRY^+^ LNd clock neurons. (A) FAFB right-hemisphere neuron visualization of nineteen unique neurons, one 5^th^ s-LNv (orange) and eighteen IPC neurons (green). The ipsilateral 5^th^ s-LNv clock neuron showed close contact points (boxed section) of 1 µm with two IPC neurons. (A’) Zoom view highlighting close points of 5^th^ s-LNv (black) and close points of IPCs (purple). Far points are shown for IPCs in green and for 5^th^ s-LNv in orange. (B) FAFB Right-hemisphere neuron visualization of twenty unique neurons two CRY^+^ LNds (red) and eighteen IPC neurons (green). Five out of eighteen IPCs showed 1 µm close points to both ipsilateral CRY^+^ LNds shown in the boxed section. (B’) Zoom view highlighting close points of CRY^+^ LNds (black) and close points of IPCs (purple). Far points are shown for IPCs in green and for CRY^+^ LNds in red. (C) MALEVNC right-hemisphere neuron visualization of eleven unique neurons, one 5^th^ s-LNv (orange) and ten IPC neurons (green). One 5^th^ s-LNv (13531) clock neuron showed close contact points (boxed section) of 1 µm with ten IPC neurons. (C’) Zoomed in image to 1 µm close points of 5^th^ s-LNv (black) and close points of IPCs (purple). Far points are shown for IPCs in green and for 5^th^ s-LNv in orange. (D) Right-hemisphere neuron visualization of twelve unique neurons two CRY^+^ LNds (red) and ten IPC neurons (green). Two CRY^+^ LNds showed 1 µm close points to ten IPCs shown in the boxed section. (D’) Zoomed in image of 1 µm close points of CRY^+^ LNd (black) and close points of IPCs (purple). Far points are shown for IPCs in green and for CRY^+^ LNds in red.

## Discussion

Brain circadian clocks across phylogeny consist of highly interconnected networks of peptidergic cells. This interconnectivity allows the central clock to maintain robust timekeeping in the face of changing internal and external states. However, central clocks must also convey time of day information outside the clock to other brain regions, a process termed clock output. A central question in the field is whether clock output requires synaptic transmission or also repurposes some of the same neuropeptides used for intra-clock communication for output signaling. Here we address this in the *Drosophila pars intercerebralis,* which has been described as a circadian output hub, with known modulation by DN1 and LNd clock neurons [27–29,31]. We demonstrate that the “core” LNv clock neurons also provide input to insulin producing cells of the PI via the classical clock neuropeptides PDF and sNPF. Functional imaging experiments confirmed that stimulation of LNvs provides time-of-day dependent input to IPCs, with strong morning activation in both male and female flies (Fig 1). Stimulation of LNvs in the presence of TTX eliminated the GCaMP response of most IPCs, however a small subset still shows an increased response, suggesting direct LNv input to some IPCs (Fig 1). Functional imaging found that *DH44^+^* neurons do not respond to LNv stimulation (Fig 2). We further find that the IPC response to PDF depends on the presence of not only PDFR but also sNPFR (Figs 3-5). Our findings also highlight the necessity of *in vivo* tools for detecting volume neurotransmission, as electron-microscopy-based connectomic approaches have limited ability to predict such interactions (Fig 6). Our data support a model in which s-LNvs provide direct and indirect input to IPCs via the neuropeptides PDF and sNPF, as well as indirect input through multisynaptic circuits (Fig 7). These multi-synaptic circuits include signaling from s-LNvs to downstream LNd and DN1p clock neurons that signal to IPCs, as previously described [29]. Neither the female FAFB nor male MCNS connectomes predict direct synaptic connections between any clock neuron and IPCs. All synaptic signaling from circadian clock neurons to IPCs thus likely requires non-clock interneurons.

**Figure 7.**
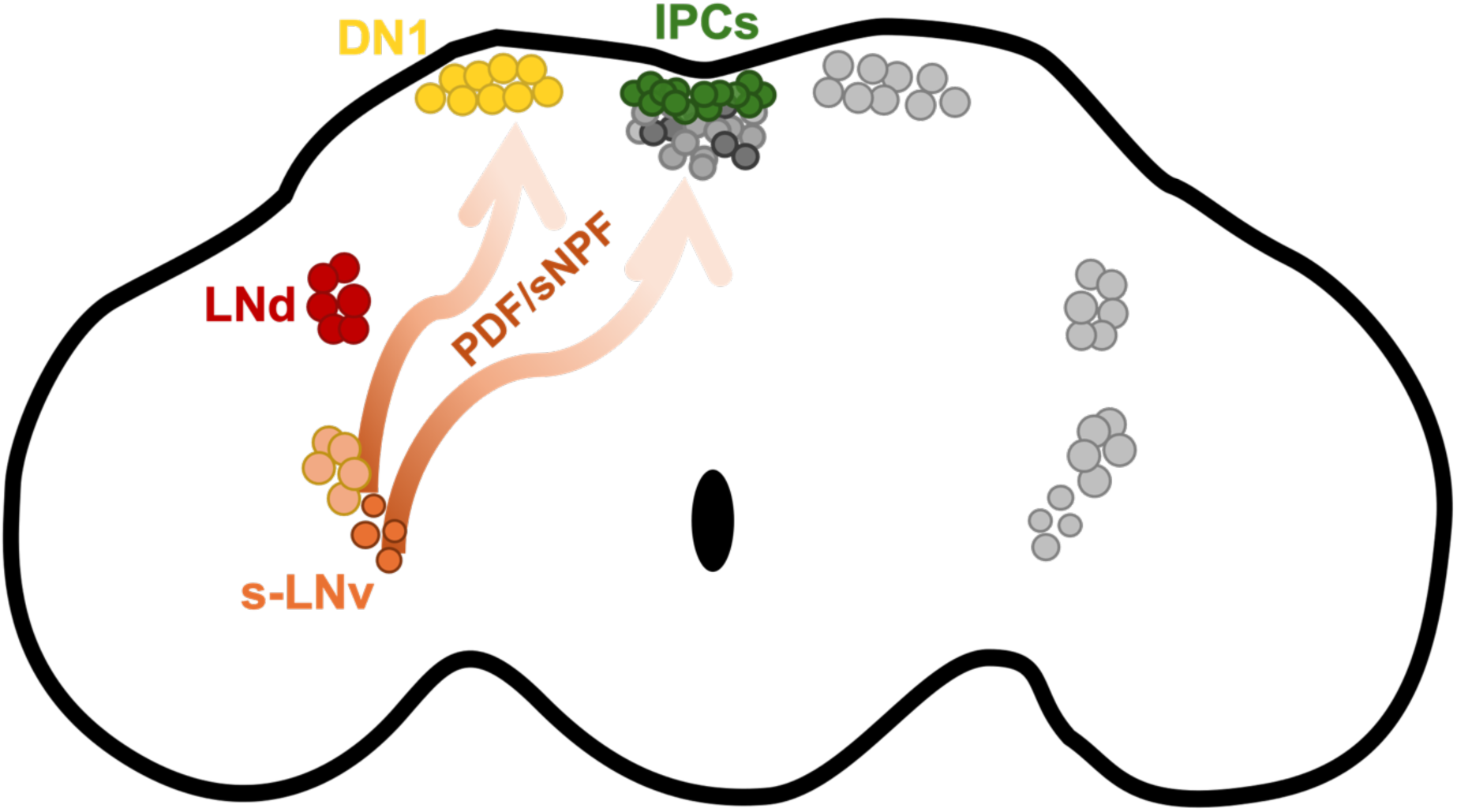
s-LNvs provide clock output signals to IPCs via peptidergic volume transmission. Small ventral lateral clock neurons (s-LNvs, orange) influence *pars intercerebralis* insulin-producing cells (IPCs, green) through both indirect and direct pathways. Indirect signaling occurs via intermediate clock neuron populations such as LNd (red) and DN1 (yellow) clock neurons that also provide input to IPCs. Synaptic inputs from s-LNvs to LNds have been identified. However, s-LNvs also signal to DN1 clock neurons and IPCs via peptidergic volume transmission of PDF and sNPF as no synaptic connections can be identified in available brain connectomes.

IPCs respond to application of sNPF and PDF with increases in cAMP even in the presence of TTX, demonstrating that IPCs can be modulated by neuropeptides expressed by the LNvs [42]. We sought to determine if LNv neurons provide direct input to IPCs by stimulating LNvs and imaging the IPC GCaMP6m response, identifying morning specific activation of IPCs by LNvs. Morning activation of IPCs upon stimulation of LNvs was comparable between sexes, although the increase in GCaMP6 signal was lower and more variable in female flies, consistent with a less dominant role for the morning oscillator in female flies (Fig 1) [47]. Both sexes showed no IPC activation upon evening LNv neuron stimulation. Morning-specific IPC activation of IPCs by LNv stimulation is aligned with a body of literature demonstrating that s-LNvs are morning-active neurons [11,19,60,61]. Ellipsoid body and s-LNv clock neurons themselves also demonstrate maximal PDF responsiveness coincident with the timing of maximal PDF release. In s-LNv clock neurons, the temporal dynamics of PDF sensitivity require the small GTPase RalA, which may control vesicle fusion dynamics to regulate PDFR isoform expression[62]. A similar mechanism could be at play in IPCs and other non-clock neurons such as those of the ellipsoid body to regulate temporal expression of GPCRs.

The average effect of morning LNv stimulation on IPC calcium levels is reduced by TTX application however a few cells per brain still appear to respond (Fig 1 A’, C’). This is consistent with previous observation of heterogenous IPC responses to clock neuron stimulation of DN1 and LNd clock neurons[28]. Recent single-cell analysis of IPCs has identified underlying heterogeneity in connectivity and gene expression that underlie this phenomenon[63]. These findings suggest that LNvs provide direct peptidergic communication to a small subset of IPCs as well as indirect modulation via multisynaptic circuits. However, it is possible that peptide release from LNv neurons themselves is impaired by TTX, as we did not directly determine that P2X2 stimulation by ATP was sufficient to drive peptide release in the presence of TTX. Consequently, we may underestimate the level of direct LNv to IPC peptide signaling.

We next sought to evaluate the roles of IPC expression of PDFR and sNPFR in the response to s-LNvs. Unfortunately, our stimulus-response paradigm requires use of two binary expression systems and five transgenes, rendering us unable to perform these experiments with IPC-specific receptor knockout. To circumvent this limitation, we turned to neuropeptide application experiments. We detected robust increases in intracellular calcium, as measured by GCaMP6m fluorescence intensity changes, upon application of both PDF and sNPF (Figs 3, 4). This is in contrast to prior findings using GCaMP3, which did not show an IPC response to PDF application in female brains [42]. This discrepancy in calcium indicator response to PDF is likely in part due to our use of a more sensitive GCaMP variant and in part due to sex and timing differences in the IPC calcium response, as shown in Fig 3. PDFR is a class B1 G-protein coupled receptor, and as with other family members seems to be primarily G_s_ coupled, leading to elevation of cAMP upon receptor activation [22–24,64]. However, PDF activation can also drive increases in intracellular calcium levels suggesting it can be G_q_ coupled [23,65]. For PDF, we tested co-application with TTX and found no significant different in IPC GCaMP response compared to PDF application alone, indicating that PDF can directly activate IPC calcium responses (Fig 3A). Direct application of PDF or sNPF peptides did not show sexually dimorphic effects on IPC-expressed GCaMP6 at any time of day, suggesting that any sexual dimorphism originates upstream of IPCs (Figs 3-5).

*DH44*^+^ neurons receive inhibitory input from LNd and DN1p clock neurons, however stimulation of LNvs did not induce changes in *DH44*^+^ neuron GCaMP6 signal in male or female flies in the morning or evening (Fig 2). Unlike IPCs, which have a peak of electrical activity and elevated calcium levels in the morning, *DH44*^+^ neurons’ activity peak is in the mid-day [29,37]. Thus, the morning-active LNvs may not be the primary drivers of daily *DH44+* neuron activity or may be involved in an indirect circuit to provide a signaling delay. Alternatively, peptidergic signaling to GPCRs in *DH44*^+^ neurons may be coupled solely to G proteins that regulate adenylate cyclase, thus failing to yield a detectable change in cytosolic calcium levels upon receptor activation.

IPC-specific CRISPR knockdown of PDFR resulted in loss of GCaMP6 response to both PDF and sNPF application, consistent with prior findings using the PDFR *Han*^5304^ mutant (Figs 3, 4) [42]. Further, IPC-specific CRISPR knockdown of sNPFR also resulted in loss of GCaMP6 response to both PDF and sNPF application (Fig 5). This could reflect compensatory changes in expression or crosstalk between the PDF and sNPF receptors necessary for maximal response to LNv derived peptides in IPCs.

There are plausible mechanisms for this cross-talk such as formation of cooperative GPCR complexes, or more likely interactions between downstream signalosome components enhancing effects on adenylate cyclase activity or calcium release [57,66–68]. A limitation of this study is the lack of direct evidence for PDFR expression in IPCs. IPC-specific sNPFR expression has been identified by both RNAseq and antibody labeling approaches, however PDFR expression has only been identified by RNAseq [38–41]. Similarly, effects of PDFR knockdown have also been found in dopaminergic neurons without molecular confirmation of protein expression in these cells [69,70]. Advances in genetic and molecular tools for probing PDFR expression and activation will be necessary to better understand the extent of PDF bulk neurotransmission.

Our findings demonstrate that LNv clock neurons morning-specifically activate IPCs, but not all PI neurons (e.g. DH44^+^ neurons). This suggests specificity of clock output signaling to the PI, with IPCs and DH44^+^ cells receiving different circadian inputs at different times of day, consistent with the different peak activity phases of these cell types[37]. We observe heterogeneity of IPC responses to LNv stimulation within the same brain, suggesting that a few IPCs may respond directly to LNv secreted signals, while most receive indirect signals through intermediary neurons. In either case, the signature clock neuropeptides, PDF and sNPF, which have primarily been implicated in intra-clock communication, also play a role in circadian clock output. In mammals, the suprachiasmatic nucleus peptide vasoactive intestinal peptide (VIP) also communicates to insulin-secreting pancreatic beta cells[71]. Thus, our data suggest that clock peptide signaling to insulin-secreting cells may be conserved from flies to humans, allowing us to further leverage the fly model to interrogate circadian control of insulin release and circulating glucose levels.

Our findings highlight the importance of *in vivo* functional experiments to identify neuromodulatory signaling by volume transmission and to probe for sparse or variable synaptic connections not detectable in connectomic datasets. Our functional connectivity, peptide application, and GPCR CRISPR knockout studies suggest that most IPCs respond to LNv activation, likely via both bulk neurotransmission and synaptic circuits containing additional intermediary neurons. A small subset may form synaptic connections, but no synaptic connections are identified in male or female fly brain connectome datasets (Fig 6). Our TTX experiments (Fig 1) do not rule out the possibility of morning-specific synaptic connections from LNvs to IPC that are not detectable in the available connectomes. Some fine neurites in the connectome are not visible or not yet traced in public datasets, which can result in underestimation of sparse synaptic connections detected experimentally. For example, prior synaptically targeted GRASP identifies sparse connections from CRY^-^ LNd clock neurons to IPCs, which are not found in connectomes [29]. Further, recent Tango-seq identified sparse synaptic connections from s-LNvs to DN1ps, also not detectable in connectomes [72]. Additionally, s-LNvs have daily remodeling of the dorsal axon terminals projecting toward the IPCs not captured by a single connectome [50,73]. However, the plastic s-LNv sites have been shown to primarily provide input to the clock, rather than output, so this remodeling may be irrelevant to LNv signaling to IPCs [74]. LNv input to IPCs is morning-specific, further work will be required to determine the molecular mechanisms underlying this temporal specificity.

Given the lack of synaptic connections from LNvs to IPCs, we turned to proximity analysis to estimate the possibility of peptidergic communication. How far any specific neuropeptide can act from its release site is not well characterized and depends on the physicochemical properties of the peptide, expression level and affinity of the receptor, and probability of degradation in transit [58,59,75,76]. Examples of peptides acting as far as hundreds of microns can be found in the literature, though it seems probable that the majority of peptide signaling is local, within less than ten microns. Proximity analysis identified PDF^+^ s-LNvs within 14.8 μm and 20 μm of IPCs in the female and male connectomes respectively (Fig 6). However, in each connectome this proximity was for a single s-LNv across both hemispheres. The responsiveness of IPCs to PDF application and to LNv stimulation implies that either PDF must be capable of volume transmission across 15-25 μm in the fly dorsal protocerebrum, or there are processes not detected in the connectome. Additional tools are necessary to unequivocally confirm PDF diffusion to IPCs, such as a PDF GPCR-activation-based (GRAB) sensor [77]. Circadian output to IPCs is quite robust, sufficient to drive diurnal IPC firing rhythms in the absence of a cell autonomous clock [28]. To accomplish this, our data suggest that intra-clock neuropeptides also serve as local clock output signals, together with multisynaptic circuits from multiple clock populations to IPCs (Fig 7).

## Materials and Methods

### Fly Stocks and Maintenance

Flies were maintained on standard cornmeal medium (BDSC) at 25° C. Flies expressing PDFR gRNA were generated using previously described protocols [53]. Briefly, gRNAs specific to the target gene PDF Receptor with FlyBase ID FBgn0260753 were identified using the web based tool for identifying targets for CRISPR/Cas9; CHOPCHOP [78]. After being flanked by tRNA and assembled and incorporated into the pCFD6 vector (Addgene), sequence-verified plasmids were sent to BestGene Inc. for injection into embryos at the attP2 and attP40 sites. See Table S1 for a list of complete genotypes and key reagents.

### GCaMP imaging of insulin producing cells

Live imaging experiments were conducted as previously described [29], with the following differences noted. Warner Instruments PM-5 perfusion chamber was used to stabilize the brain under nylon fibers attached to a platinum wire frame. GCaMP6m calcium imaging was performed on a Leica TCS SP8 confocal microscope. Z-stacks were acquired every 11 s when GCaMP6m was expressed in *DH44*^+^ cells and 14 s in *DILP2*^+^ cells. Constant perfusion of artificial hemolymph HL3 [45] was done for the duration of imaging session, with addition of stimulants after a baseline reading of 5 z-stacks. ATP was used at 2.5 mM in HL3 at pH 7.1 to activate P2X2. Tetrodotoxin (TTX, VWR 89160-628) was used at 2 μM in HL3 and ATP. 10 μM of PDF (*NSELINSLLSLPKNMNDAa,* GenScript) or sNPF (*AQRSPSLRLRFa,* GenScript) in 0.1% DMSO in HL3 was used for neuropeptidergic activation experiments, and 0.1% DMSO in HL3 was used as a vehicle control in peptide application experiments. Stimulation readings were done for 15 stacks after baseline. Images were acquired using a Leica TCS SP8 laser scanning confocal microscope with a 40x/1.3 NA and a 1-3 μm z-step size.

Image processing and measurement of fluorescence intensity was performed in FIJI as previously described [79]. Briefly, for each time point, z-stacks were converted into sum-intensity projections, which were subsequently used for all analyses. To compensate for any minor x-y drift during imaging, the FIJI Image Stabilizer plugin was applied to the projected image sequences. Individual GCaMP-expressing cell bodies were manually outlined as regions of interest (ROIs) and the mean fluorescence intensity within each ROI was quantified for each z-stack over time. Fluorescence responses were normalized according to the formula ΔF/F = (F_n_ − F_0_)/F_0_, where F_n_ represents the fluorescence intensity at a given time point and F_0_ corresponds to the average fluorescence measured during the five baseline frames immediately preceding ATP application.

Statistical analysis and creation of graphs was performed in OriginLab 2025b. Peak calcium responses for individual cells were determined by calculating the difference between the highest ΔF/F value observed during ATP/synthetic peptide/vehicle exposure and the mean ΔF/F during the five baseline frames before ATP/synthetic peptide/vehicle delivery. Brains exhibiting unstable baseline fluorescence were excluded from further analysis. All cells within a condition (by genotype, sex, circadian time, perfusate application) were averaged to create ΔF/F time course plots. Although ΔF/F data were determined for each cell, cell-level data were averaged by brain, as each brain represents an independent sample for analysis of variance (ANOVA). ΔF/F values were tested for normality by genotype for each condition using the Shapiro-Wilk normality test. Statistical comparisons of maximum and minimum ΔF/F values by brain among experimental groups were performed using one-way ANOVA followed by Tukey’s multiple-comparisons *post hoc* test for normally distributed data and Kruskal-Wallis ANOVA for non-normal data.

### Connectomic analysis

We first used cell type, super class, and neurotransmitter prediction type data sets from FlyWire Brain Dataset Female Adult Fly Brain (FAFB version 783) and the male CNS (MCNS version 0.9) to build a master classification table in R studio for each neuron root id. Each neuron root with its metadata was then matched to its respective neuron skeleton. Neuron skeletons for the eighteen IPC cells, two 5^th^-s-LNvs, and four CRY^+^ LNds across both brain hemispheres were used for our proximity analysis. For the analysis, we ran a python KD-tree distance analysis [5]. In the proximity analysis, we adjusted the distance analysis to 1 µm from 15 µm (in FAFB) or 20 µm (in MCNS) and then visualized the right hemisphere. We also visualized a zoomed image for each close point.

## Acknowledgements

We are grateful to Meet Zandawala and Zeyu (Ray) Chang for training and assistance in connectomic analysis. We thank Maria de la Paz Fernandez, Meet Zandawala, and all members of the Barber Lab for helpful discussions and comments on the manuscript. We acknowledge the Waksman Institute Shared Imaging Facility at Rutgers, The State University of New Jersey for confocal microscope use and support. Stocks obtained from the Bloomington Drosophila Stock Center (NIH P40OD018537) were used in this study.

