## Supplementary Information for "Drosophila core circadian clock neurons peptidergically regulate activity of insulin-producing cells"

### Hameed et al. Supporting Information

**Video S1. *Ex vivo* live imaging of the *Drosophila pars intercerebralis***

DILP2+ neurons are labeled with GCaMP6m (blue), which is expressed throughout the cell, and nuclear-localized mCherry (magenta). The scale bar represents 20 μm length and is shown throughout the video. ATP was applied following the baseline recording, as indicated. Images were acquired on a Leica SP8 confocal microscope at a resolution of 256 × 256 pixels. Z-stack images were projected into a single plane using maximum-intensity projection.

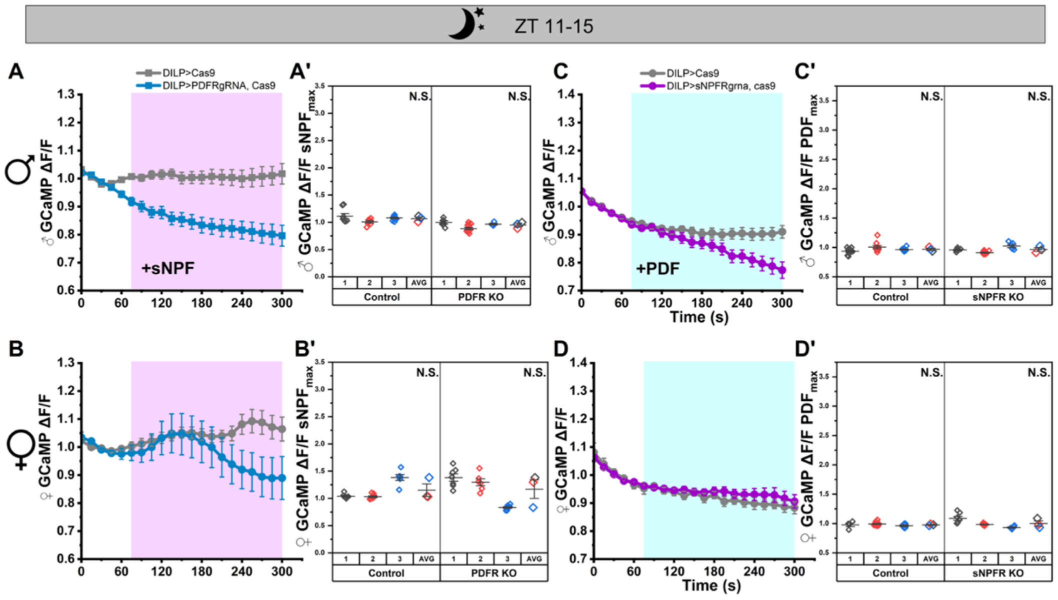


**Fig S2. IPCs maintain lack of evening stimulation by LNv neuropeptides upon knockout of non-cognate receptors.**

(A) GCaMP6m signal over time in IPCs with (blue: n=3 brains, 21 cells) and without (gray: n= 3 brains, 26 cells) PDFR knockout from male brains imaged at ZT 11-15 during perfusion of artificial hemolymph, followed by perfusion of 10 μM sNPF. Magenta area indicates timing of sNPF application. Data represented as mean ± SEM. (A') Maximum change in GCaMP intensity during activation for each cell by brain and brain averages by genetic line from panel A. Each small point represents an individual cell, while larger symbols at the left of each line panel indicate the mean response per brain matching individual symbol colors by brain (B') Plotted as in panel A' for panel B. (C) GCaMP6m signal over time in IPCs with (magenta: n=3 brains, 32 cells) and without (gray: n=4 brains, 34 cells) sNPFR knockout from male brains imaged at ZT 11-15 during perfusion of artificial hemolymph, followed by perfusion of 10 μM PDF. Blue area indicates timing of PDF application. Data represented as mean ± SEM. (C') Maximum change in GCaMP during PDF application for each cell from panel C. Each point is an individual cell. (D) Plotted as in panel C from female brains (magenta: n=3 brains, 21 cells, gray: n= 3 brains, 23 cells). (D') Plotted as in panel C' for panel D. Data are presented as mean ± SEM for cells. Brain means that do not share a letter are significantly different; N.S. indicates no significant difference. One-way ANOVA analysis with Tukey's multiple comparisons done across genetic lines. $P \leq 0.05$

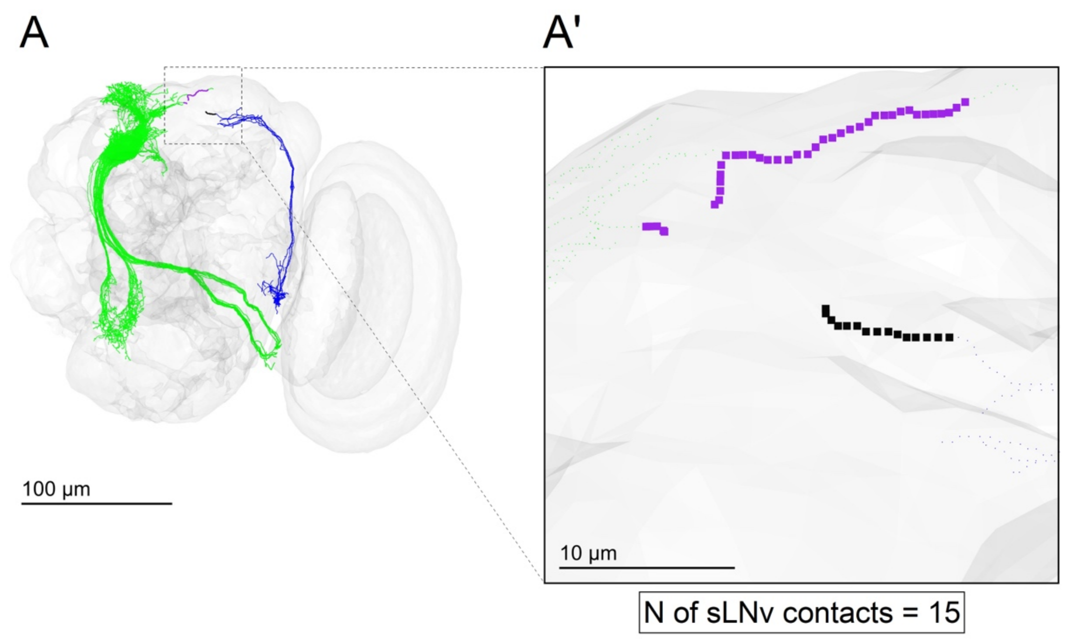


**Fig. S3. A single male s-LNv comes within 25 µm of any IPC**.

(A) MALEVNC right-hemisphere neuron visualization of two unique neurons, one s-LNv (blue) and one IPC neuron (green). The s-LNv clock neuron showed fifteen close contact points (boxed section) of 25 µm with one IPC neuron. (A') Zoomed in image to 25 µm close points of s-LNv (black) and close points of IPCs (purple). Far points are shown for IPCs in green and for s-LNv in blue.

**Table S1.** Key Reagents

| REAGENT or RESOURCE | SOURCE | IDENTIFIER |
|---|---|---|
| **Antibodies** | | |
| Mouse anti-rat CD2 | Genetex | Cat# GTX75123 |
| Rat monoclonal anti-RFP | ChromoTek | Cat# 5f8-100, RRID: AB_2336064 |
| Cy3-AffiniPure Donkey Anti-Rat IgG (H+L) | Jackson ImmunoResearch Labs | Cat# 712-165-153, RRID: AB_2340667 |
| Cy5-AffiniPure Donkey Anti-Mouse IgG (H+L | Jackson ImmunoResearch Labs | Cat# 715-175-151, RRID: AB_2340820 |
| **Chemicals, Peptides, and Recombinant Proteins** | | |
| PDF peptide (NSELINSLLSLPKNMNDAa) | GenScript | Custom synthesis |
| sNPF peptide (AQRSPSLRLRFa) | GenScript | Custom synthesis |
| Adenosine 5'-triphosphate disodium salt hydrate | Millipore Sigma | Cat#: A26209-10G, CAS: 34369-07-8 |
| Tetrodotoxin | VWR | 89160-628 |
| **Fly Lines** | | |
| *D. melanogaster*: *Dh44-GAL4 (attp2)* | VDRC | VDRC: 207474 Flybase: FBti0169412 |
| *D. melanogaster: Dh44-LexA (attp40)* | BDSC | RRID: BDSC_53591 |
| *D. melanogaster:* DILP2-Gal4 | BDSC | RRID:BDSC_37516 |
| D. melanogaster:PDF-LexA | BDSC | RRID:BDSC_52523 |
| *D. melanogaster:* LexAop-P2X2 | BDSC | RRID:BDSC_76030 |
| *D. melanogaster: 20X UAS-IVS-GCaMP6m* | BDSC | RRID:BDSC_42750 |
| *D. melanogaster: Iso31* | Ryder *et al.* Genetics 2004 | RRID:BDSC_5905 |
| *D. melanogaster: InsP3-Gal4* | Gift from Dr. Michael Pankratz | Buch *et al.* Cell Metabolism 2008 |
| *D. melanogaster: UAS-Cas9.P2* | BDSC | RRID BDSC 58986 |
| D. melanogaster: PDFRgRNA (attP2) | This Study | |
| *D. melanogaster: sNPFRgRNA* | Schlichting et al. Proc Nat Acad Sci USA 2022 | RRID:BDSC_97854 |

| | | |
|---|---|---|
| *D. melanogaster: UAS-mCherry.nls* | BDSC | RRID:BDSC 38425 |
| **Fly line annotations** | | |
| *>DILP | ;DILP2-Gal4/UAS-mcherry.nls, UAS-GCaMP6m; TM2, ubx/Aop-P2X2 | |
| PDF>DILP | ;DILP2-Gal4/UAS-mcherry.nls, UAS-GCaMP6m; PDF-LexA/Aop-P2X2 | |
| *>DH44 | ;DH44-Gal4/UAS-mcherry.nls, UAS-GCaMP6m; TM2, ubx/Aop-P2X2 | |
| PDF>DH44 | ;DH44-Gal4/UAS-mcherry.nls, UAS-GCaMP6m; PDF-LexA/Aop-P2X2 | |
| DILP>cas9 | ;DILP2-Gal4/Cyo; UAS-Cas9.P2/TM6, sb | |
| DILP>PDFR gRNA, cas9 | ;DILP2-Gal4/UAS-PDFRgRNA; UAS-Cas9.P2/TM6, sb | |
| DILP>sNPFRgRNA, cas9 | ;DILP2-Gal4/UAS-sNPFRgRNA; UAS-Cas9.P2/TM6, sb | |
| **Primers and Plasmids** | | |
| PCFD6 plasmid | Addgene | Cat# 73915 |
| PDFR guide 1: CGGCCCGGGTTCGATTCCCGGCCGATGCACTCCCACATTCACAGCAAGGAGGGTTTCAGAGCTATGCTGGAAAC | | |
| PDFR guide 2: AAATGTGCAGAAGATCAGCAGGGGTTTCAGAGCTATGCTGGAAAC | | |
| PDFR guide 3: CTGTCGAACATTCTCGACTGCGGGTTTCAGAGCTATGCTGGAAAC | | |
| PDFR mRNA Primer Pair 1: GCGTTCGTTTCGCTTTTCCA / CAAACTCGCGTTGGAAGCAC | | |
| PDFR mRNA Primer Pair 2: TGGCGTTCGTTTCGCTTTTC / CGGGTTGAAATCAATTGGGCA | | |
| PDFR mRNA Primer Pair 3: TCATAACGGCTTCGGCACTC / CGGGTTGAAATCAATTGGGCA | | |
| **Software** | | |
| FIJI distribution of ImageJ | Schindelin *et al.* Nat Methods 2012 | https://fiji.sc |
| OriginPro Version 2020 | OriginLab Corporation, Northampton, MA | |
| CHOPCHOP | Labun et al. Nucleic Acids Res 2019 | https://chopchop.cbu.uib.no/ |

**Table S1: Key Resources.** This table provides all antibodies, reagents, peptides, fly lines, plasmids, primers, full fly genotypes and software distributions used to facilitate reproducibility of results.
